# Cryo-EM structure of the conjugation H-pilus reveals the cyclic nature of the TrhA pilin

**DOI:** 10.1101/2024.12.30.630807

**Authors:** Naito Ishimoto, Joshua L.C. Wong, Nanki Singh, Sally Shirran, Shan He, Chloe Seddon, Olivia Wright-Paramio, Carlos Balsalobre, Ravi R. Sonani, Abigail Clements, Edward H. Egelman, Gad Frankel, Konstantinos Beis

**Affiliations:** Rutherford Appleton Laboratory, Research Complex at Harwell, Didcot, Oxfordshire, UK; Department of Life Sciences, Imperial College London, London, UK; BSRC Mass Spectrometry & Proteomics Facility, University of St Andrews, St Andrews, UK; Department de Genètica, Microbiologia i Estadística, Facultat de Biologia. Universitat de Barcelona, Barcelona, Spain; Department of Biochemistry and Molecular Genetics, University of Virginia, Charlottesville, VA, USA

**Keywords:** conjugation, conjugation pilus, mating pair formation, cyclization, mass spectrometry

## Abstract

Conjugation, the major driver of the spread of antimicrobial resistance genes, relies on a conjugation pilus for DNA transfer. Conjugative pili, such as the F-pilus, are dynamic tubular structures, composed of a polymerized pilin, that mediate the initial donor–recipient interactions, a process known as mating pair formation (MPF). IncH are low-copy-number plasmids, traditionally considered broad host range, which are found in bacteria infecting both humans and animals. The reference IncHI1 plasmid R27, isolated from *Salmonella enterica* serovar Typhi, encodes the conjugative H-pilus subunit TrhA containing 74 residues after cleavage of the signal sequence. Here, we show that the H-pilus forms long filamentous structures that mediate MPF, and describe its cryo electron-microscopic (cryo-EM) structure at 2.2 Å resolution. Like the F pilus, the H-pilin subunits form helical assemblies with phospholipid molecules at a stochiometric ratio of 1:1. While there were previous reports that the T-pilus from *Agrobacterium tumefaciens* was composed of cyclic subunits, three recent cryo-EM structures of the T-pilus found no such cyclization. Here, we report that the H-pilin is cyclic, with a covalent bond connecting the peptide backbone between the N- and C-termini. Both the cryo-EM map and mass spectrometry revealed cleavage of the last five residues of the pilin, followed by cyclization via condensation of the amine and carboxylate residues. The cyclic nature of the pilin could stabilize the pilus and may explain the high incidence of IncH plasmid dissemination.

**Significance:** A major medical challenge is the spread of bacteria which are resistant to antibiotics. The resistance genes are spread via mobilized DNA, mainly via a process named conjugation. During conjugation, a resistant bacterium (donor), transfers the resistance DNA to another bacterium (recipient) in a contact-dependent manner. The initial donor-recipient interaction is mediated by a hollow filament expressed by the donor, named the conjugation pilus, that binds the recipient. This pilus is built via polymerization of a small protein subunit, pilin. Here, we report the atomic structure of the H-pilus, whose pilin subunit has an unusual cyclic structure where the N- and C-termini of the protein are covalently linked by a peptide bond.

## Introduction

Conjugation is a major mechanism of horizontal gene transfer, enabling the exchange of genetic material between bacterial cells. This process is central to bacterial evolution, particularly in exchange of virulence and metabolic genes as well as in the dissemination of antibiotic resistance genes. During conjugation plasmid DNA is transferred unidirectionally from one bacterium (donor) to another (recipient) in a contact-dependent manner (reviewed in (1)). DNA transfer is facilitated by a type IV secretion system (T4SS) nanomachine, embedded in the bacterial cell wall, which is extended by a long hollow filament, known as the conjugation, mating, or sex pilus. Functionally, the pilus is involved in mating pair formation (MPF), linking donors and recipients at a distance (1). In IncF plasmids, MPF has been shown to enable low efficiency DNA transfer via the hollow conjugation pilus (2),(3). This is followed by pilus retraction, which enable donor and recipient bacteria to form mating aggregates via intimate interactions between the plasmid encoded outer membrane protein TraN in the donor and an outer membrane protein (OMP) receptor in the recipient, a process known as mating pair stabilization (MPS) (4). We have recently shown the existence of at least four distinct TraN isoforms, each binding a distinct OMP receptor and influencing conjugation species specificity (5),(6). By stabilizing donor-recipient interactions, MPS enables efficient DNA transfer.

While pili were originally defined as all cell surface filaments that are not flagella, it has become clear that there are many different classes of pili that have evolved independently. Beyond conjugation, pili perform important other roles. For instance, type 1 pili in uropathogenic *Escherichia coli* (UPEC) mediate adhesion to host cells, enabling colonization and infection (7). Type III pili (e.g. the *Klebsiella pneumoniae* Mrk) are involved in biofilm formation (8). Type IV pili contribute to both adhesion and motility via a process called twitching motility, allowing cells to glide across surfaces (9), as well as functioning as competence pili that mediate DNA uptake in natural transformation (10). Pili also assist in immune evasion by enabling adherence to immune-inaccessible niches (11). In addition, pili can undergo antigenic variation, which helps pathogens avoid detection by the host immune system (12). Pili also mediate DNA transfer across kingdoms, for example the transfer of the Ti plasmid via the T-pilus from *Agrobacterium tumefaciens* to plant cells (13).

Pilins that polymerize to form extracellular pili are synthesized in the cytoplasm as precursors with a leader peptide that directs their export to the periplasm via the Sec pathway, where the leader peptide is cleaved to produce the mature pilins (14). Structures of mating or conjugation pili from the IncF and IncW incompatibility plasmids have revealed that the pilins polymerize into a helical arrangement with tightly bound phospholipid molecules, forming thin, tubular filaments ∼8 nanometers in diameter that can extend several micrometers in length (15), (16). Conjugation pilins are composed of three α helices, α1-3. Binding of the phospholipid is coordinated by α3 through hydrophobic interactions with the acyl chain of the phospholipid (15). Recently, it has been shown based upon structural analysis that conjugation pili in archaea are homologs of the bacterial ones (17).

IncH plasmids are divided into IncHI and IncHII, and the former is further divided into IncHI1, IncHI2 and IncHI3 (18). IncH plasmids, particularly IncHI1 and IncHI2 subtypes, are frequently associated with resistance to a wide range of antibiotics, including aminoglycosides, beta-lactams, sulfonamides, tetracyclines, chloramphenicol, and macrolides. IncH plasmids have been identified in several clinically significant Gram-negative bacterial pathogens (e.g. *K, pneumoniae* and *Acinetobacter baumannii*), where they play a central role in mediating multidrug resistance (MDR). In recent years IncH plasmids have been shown to play a major role in the spread of New Delhi metallo-β-lactamase-1 (NDM-1) (18). IncH plasmids harbor a conjugation system resembling the *tra* operon found in IncF plasmids, which fall under the MPFF classification (19). The key feature of the IncH plasmids is that conjugation rate is optimal at temperatures between 22 and 30°C and severely drops at higher temperatures (20), (21). This feature suggests that the dissemination of resistances by IncHI plasmids is promoted in water and soil environments. Accordingly it has been shown that expression of conjugative machinery genes is expressed al low temperatures (25°C) and repressed at high temperatures (37°C) (20), (21).

Using a derivative of the reference IncHI1 plasmid, R27, a tetracycline resistance plasmid isolated from *Salmonella enterica* serovar Typhi (20), we determined the cryo-EM structure of the H-pilus at 2.2 Å resolution. Like the F (15) and W (16) pili, the H-pilin subunits, TrhA, form helical assemblies with phospholipid molecules at a stochiometric ratio of 1:1. Uniquely, we found that unlike the F and W pilin structures, TrhA is cyclic, formed by a covalent bond connecting the peptide backbone between the N- and C-termini. Mass spectrometry data further validated the structure and confirmed that the last five residues of the mature pilin are cleaved, followed by cyclization via condensation of the amine and carboxylate residues. The cyclic nature of TrhA may provide the H-pilus with greater stability, which could potentially enable more efficient plasmid transfer during MPF.

## Results

### Purification of the H-pilus

In order to optimize expression conditions of the derepressed *htdA* R27 mutant (drR27 in which resistance was swapped from tetracycline to chloramphenicol) (21), which constitutively expresses the conjugation operons at 25°C, we first conducted conjugation assays using a tryptophan auxotrophic *E. coli* donor and wild type (WT) *E. coli*, *Citrobacter rodentium* and *K. pneumoniae* recipients. Briefly, overnight donor and recipient cultures were mixed in a ratio of 10:1, diluted 1 in 25 in PBS and 40 μl spotted onto LB agar plates. Following incubation at 37°C for 3 h, conjugation was allowed to proceed overnight at 25 °C. Transconjugates were selected by plating the conjugation mixture on M9 minimal media plates containing chloramphenicol. This revealed that almost 100% of the *E. coli* and *C. rodentium* recipients were transconjugates, while conjugation into *K. pneumoniae* was less efficient (Fig. 1A).

**Figure 1.**
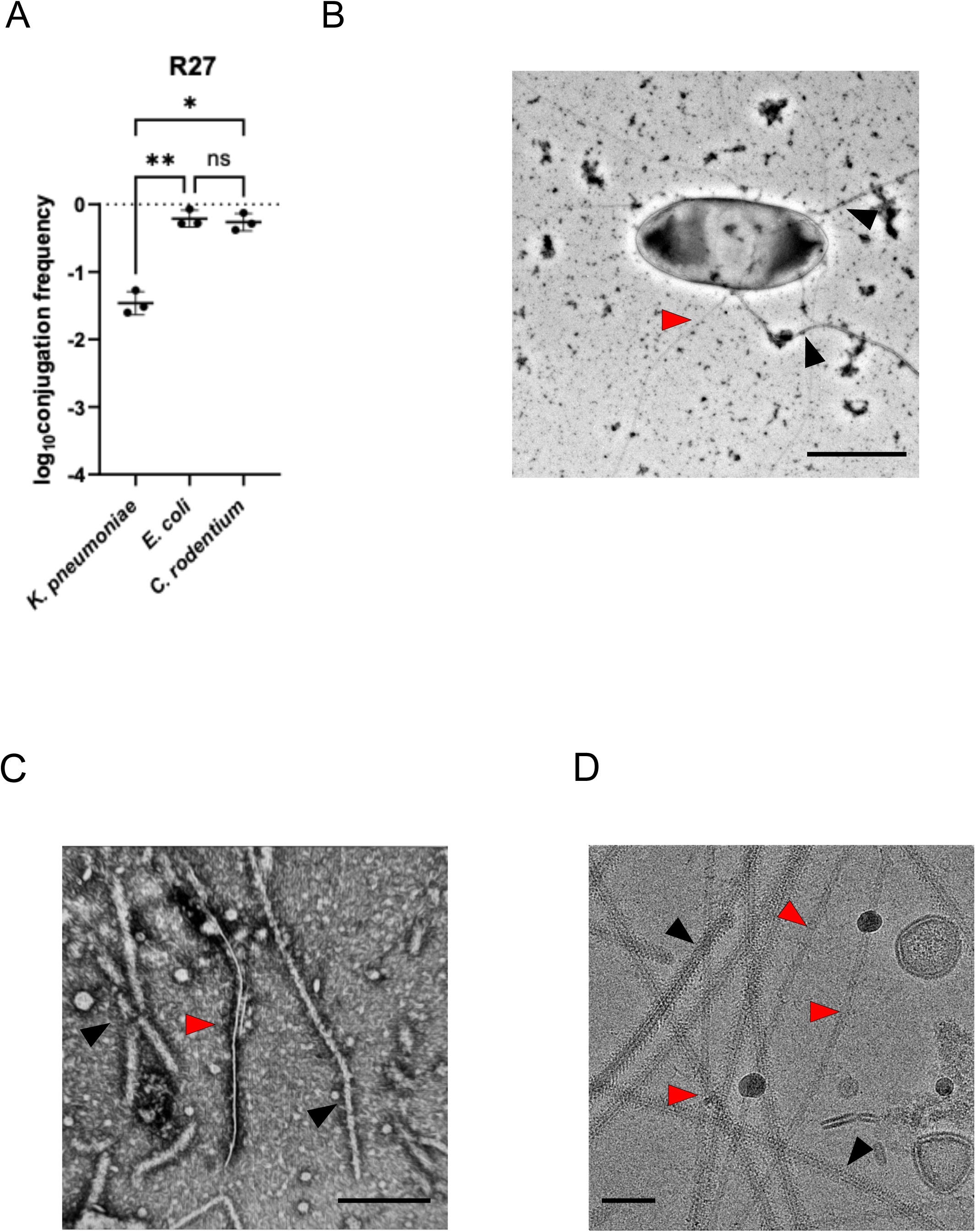
Purification of the R27 H-pilus. (A) Conjugation of R27 from *E. coli* MG1655 Δ*trp* donor into *K. pneumoniae, E. coli* or *C. rodentium* recipients. Data were statistically analyzed by a two paired *t*-test. Statistical significance is marked by * (p < 0.05), ** (p < 0.01) and ns (not significant). Data presented are representative of three biological repeats with the individual repeats and the overall average for each data set being shown. (B) Negative stain EM of MG1655 Δ*fimA* cells carrying the drR27, expressing the H-pilus (red arrows); native MG1655 flagellar (black arrows) are also visible. Scale bar is 1 μm. (C) The negative stain EM of the purified H-pilus shows the presence of both H-pilus (red arrow) and flagella (black arrows). The scale bar is 200 nm. (D) Under cryo-EM imaging, H-pilus (red arrows) appears as a hollow featureless structure whereas the flagella (black arrows) display more internal structure. The scale bar is 50 nm.

To visualize the H-pilus, we transformed drR27 into *E. coli* MG1655 Δ*fimA*, lacking expression of type I pili. Following culturing at 25°C, we used negative stain EM to visualize surface appendages. We detected both thin pilus structures of ∼ 90 Å diameter, presumably the H pili, and filamentous structures with a diameter of around 200 Å that are flagella (Fig. 1B). Finally, we used similar growth conduction to purify the H-pilus. In brief, we passed the cells through 25 G needles to shave the filaments which were precipitated by PEG 6,000 and dialyzed in imaging buffer (see Materials and Methods). The purified filaments were deposited onto carbon coasted grids and stained by uranyl acetate. The negative stain EM of the purified samples show filaments corresponding to flagella and H-pili (Fig. 1C). Cryo-EM imaging of the purified sample further confirmed the presence of flagella and H-pili (Fig. 1D). In raw images flagella showed structural features whereas the H-pili appeared as hollow tubes.

### Cryo-EM structure of the H-pilus

We processed data using standard helical reconstruction in cryoSPARC and reconstructed a map of the H-pilus at 2.2 Å resolution (map:map FSC, 0.143 threshold), by imposing helical (rise 12.14 Å, twist 28.98°) and C5 symmetries (22) (Fig. 2A,B; Supp. Fig.1). The 5-fold rotational symmetry is consistent with other characterized pili, including F and F-like pili (15), (17), while the pKpQIL pilus does not have any rotational symmetry (23). The assembled pilus structure displays an outer diameter of ∼ 90 Å and a lumen diameter of ∼ 22 Å, which is similar to previously reported conjugation pilus structures (Fig. 2B). The high-resolution density map has allowed us to build the atomic model of the H-pilus for both the main chain and side chains (Fig. 2C,D). Our analysis revealed clear phospholipid density in the map, indicating a precise 1:1 stoichiometry between pilus subunits and phospholipids. Although we have not identified the exact lipid, the density map clearly delineated the branched hydrocarbon chains, which were fit as PG32:1. The phosphate groups in the head region showed some flexibility, resulting in weaker density, but the overall lipid architecture was well-defined (Fig. 2C). Additionally, we identified and modelled several bound water molecules per subunit, which are located on both the inner and outer surfaces of the pilus (Fig. 2C,D; Supp. Fig. 2).

**Figure 2.**
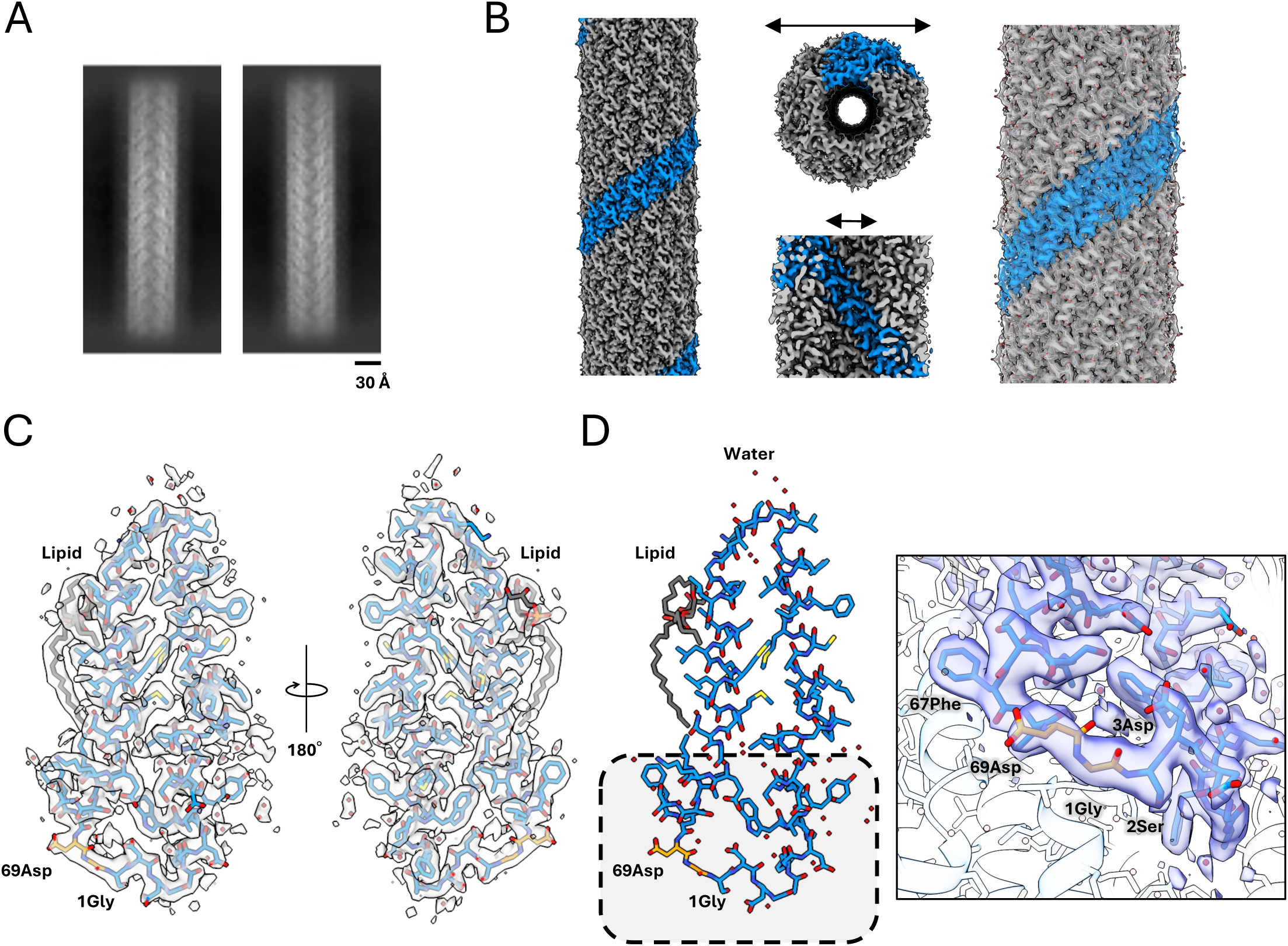
Cryo-EM structure of H-pilus. (A) The 2D class averages of H-pilus show high resolution structural features. The scale bar is 20 Å. (B) Cryo-EM map of H-pilus. (Left) One right-handed 5-start strand is shown in blue. The others are shown in grey. (Middle) The H-pilus has an outer diameter of ∼90 Å and an inner lumen diameter of ∼22 Å. (Right) Several TrhA pilin atomic models have been built within the map and are shown as sticks. (C) The high-resolution map shows density for the side chains including the site of cyclization. All the residues have been fit into the map. The TrhA pilin is shown in blue sticks within the cryo-EM map. The PG lipid is shown in grey. Water molecules are shown as red dots. (D) Structural model of the TrhA pilin, indicating the site of N- to C-terminus cyclization (boxed region). The cyclization of Gly1 to Asn69 is shown in orange sticks. Close up view of the cryo-EM map indicating the site of cyclization.

### The TrhA pilin is cyclized

Analysis of the H-pilus structure revealed cyclization between the N- and C-termini of the TrhA pilin. The density map unambiguously demonstrated the formation of a peptide bond between Gly1 and Asn69 (Fig. 2C; Supp. Fig. 2). Although the mature TrhA pilin consists of 74 amino acids, only density for the first 69 residues was visible in the maps, indicating that the five C-terminal residues were cleaved during the cyclization process. As the H-pilus was resistant to tryptic digest studies, we validated the structural observation by performing electrospray ionisation mass spectrometry (ESI-MS). While the expected molecular weight for the mature TrhA is 7593.8 Da, the experimental mass was determined to be 7124.5 Da, which precisely corresponds to the loss of A_70_GIPD_74_ (451.3Da) and the theoretical mass of residues 1-69 (7142.5 Da) minus 18 Da, representing the loss of one water molecule during the condensation of the Gly1 amine and Asn69 carboxylate residues as a result of the peptide bond formation during cyclization (Fig. 3). The unambiguous density for the cyclization site, coupled with the mass spectrometry results, definitively establishes the cleavage of five C-terminal residues in the mature H-pilus structure.

**Figure 3.**
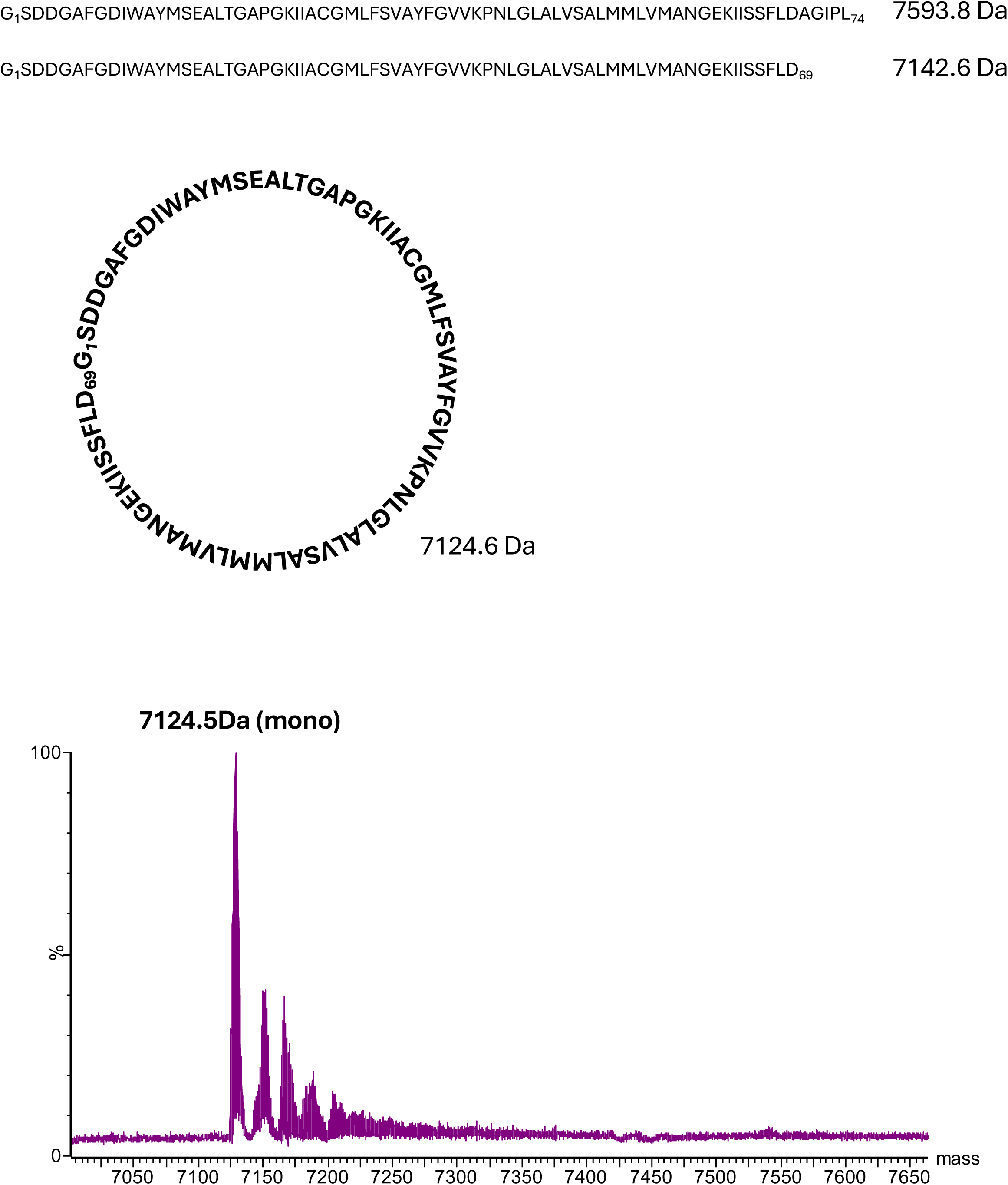
Mass spectrometry analysis of the purified TrhA pilin. The predicted molecular weight for the mature ThrA is 7593.8 Da. The measured molecular weight was 7124.5 Da, which corresponds to the loss of the five terminal residues and subsequent cyclization with the loss of a water molecule (18 Da). This value agrees with the theoretical molecular weight of the cyclic pilin at 7124.6 Da. Acyclic TrhA with the loss of the terminal residues has a predicted molecular weight of 7142.6 Da.

### H-pilus assembly

Like other pilins, the TrhA pilin is comprised of three α-helices: α1 (Gly8-Thr19), α2 (Ala21-Phe37), and α3 (Gly45-Phe66), connected by short loops (Fig. 4A). Structural comparison shows distinct structural features with six previously characterized pili: pED208 (PDB 5LEG)(15), F (PDB 9MOQ), pKpQIL (PDB 7JSV)(23), T (PDB 8EXH)(17), pKM101 (PDB 8CW4)(24), and R388 (PDB 8S6H)(16).The pED208, F, and pKpQIL-pilus structures have straight α3 helices with loops between α1-α2, whereas pKM101, T, and R388-pilus structures have kinked α3 helices with straight α1-α2 regions (Fig. 4B). The H-pilus structure uniquely combines elements from both structural groups, featuring a moderately kinked α3 helix with less a pronounced kink than in other structures, while maintaining an α1-α2 connecting loop (Fig. 4C). The loop that connects α1 and α2 in H-pilus is less kinked and more structured with an additional turn, similar to the α1/2 (fused α1 and α2 helices) of the T pilus, pKM101 and R388. The distinctive kink angle in the H-pilus, which is smaller than in other pili, appears to be directly related to its cyclization. The formation of a cyclic structure between N- and C-termini brings the α1-2 and α3 helical regions closer, likely contributing to the reduced kink angle. The structure of the pilus lumen reveals a helix-loop-helix arrangement between α2 and α3, with the loop region showing a positive charge oriented toward the lumen (Fig. 4A,C; Supp. Fig. 3). The surface electrostatic potential for the lumen will depend upon the head group of the bound lipid. For pED208, it was shown that the lumen would have a positive potential without the lipid, but the PG headgroups changed the overall potential to negative (15). For the T-pilus, the potential of the lumen was positive with and without the inclusion of the PE headgroup (17). Given that we have not unambiguously identified the bound lipid, we do not know the overall surface potential of the lumen. The exterior surface of the pilus exhibits hydrophilic characteristics, featuring extensive water molecule associations and a pronounced positive charge distribution (Figs. 2D, 5A; Supp. Figs. 3,4). The Lys24 side chain within the cyclized region, (Fig. 5A), likely enhances subunit interactions, thereby contributing to the structural stability of the assembled pilus.

**Figure 4.**
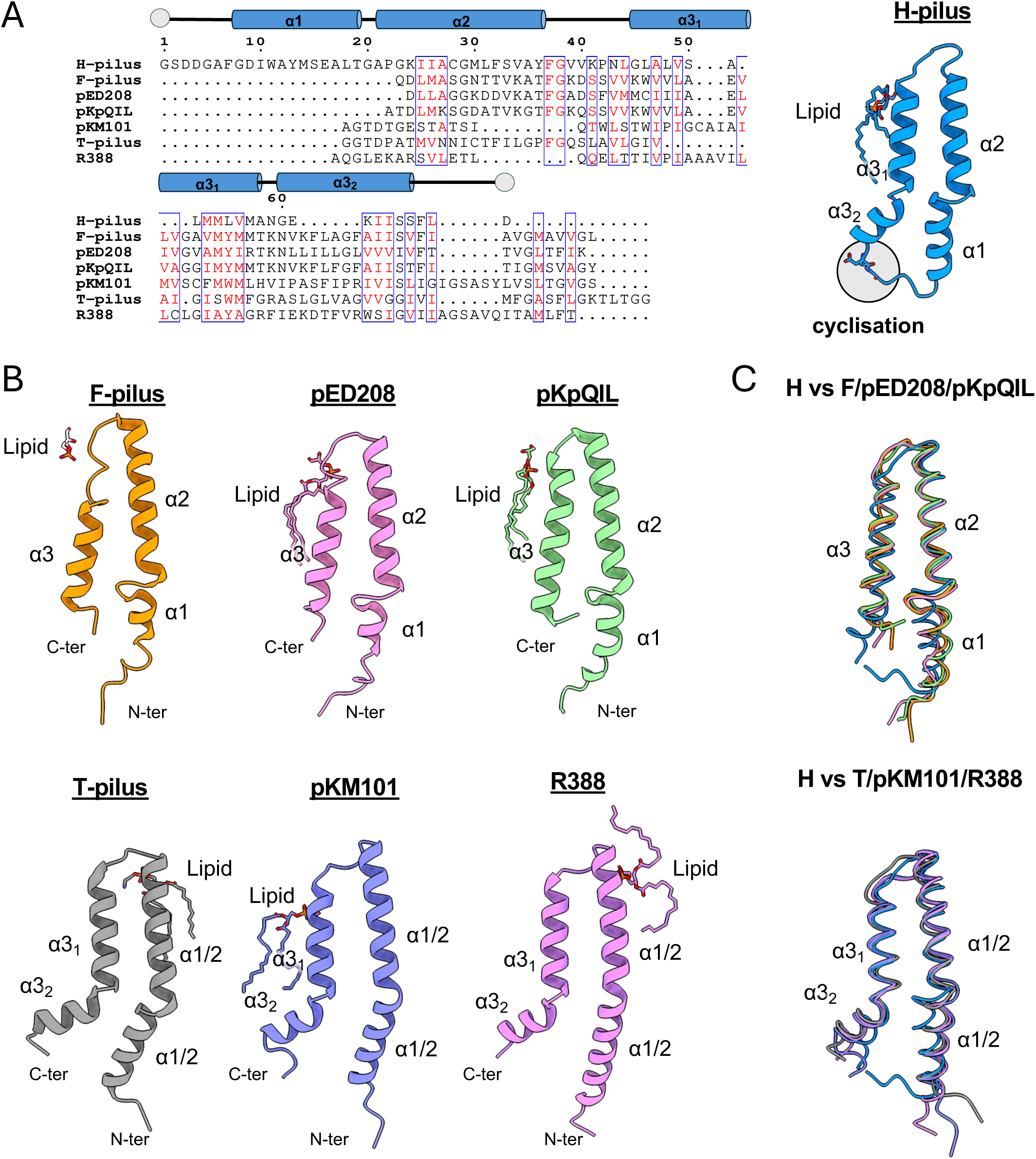
Comparison of H-pilus with other mating pilus structures. (A) Sequence alignment of mature pilins. The tube shows the location of the α-helices of the TrhA. N and C termini are shown in grey circles; similar residues are shown in red, and similar characteristic residues are boxed. The alignment was performed in CLUSTALW. The structure of the TrhA pilin is shown in cartoon. (B) The pilin structures of known pilins bound to their lipid are shown in cartoon: F-pilus (9MOQ), pED208 (5LEG), pKpQIL (7JSV), T-pilus (8EXH), pKM101 (8CW4), R388 (8S6H). The main structural difference between the top and bottom rows is the kink of α3. (C) Superposition of TrhA with the individual pilins from panel (B). TrhA shares structural similarities from both conformational classes.

**Figure 5.**
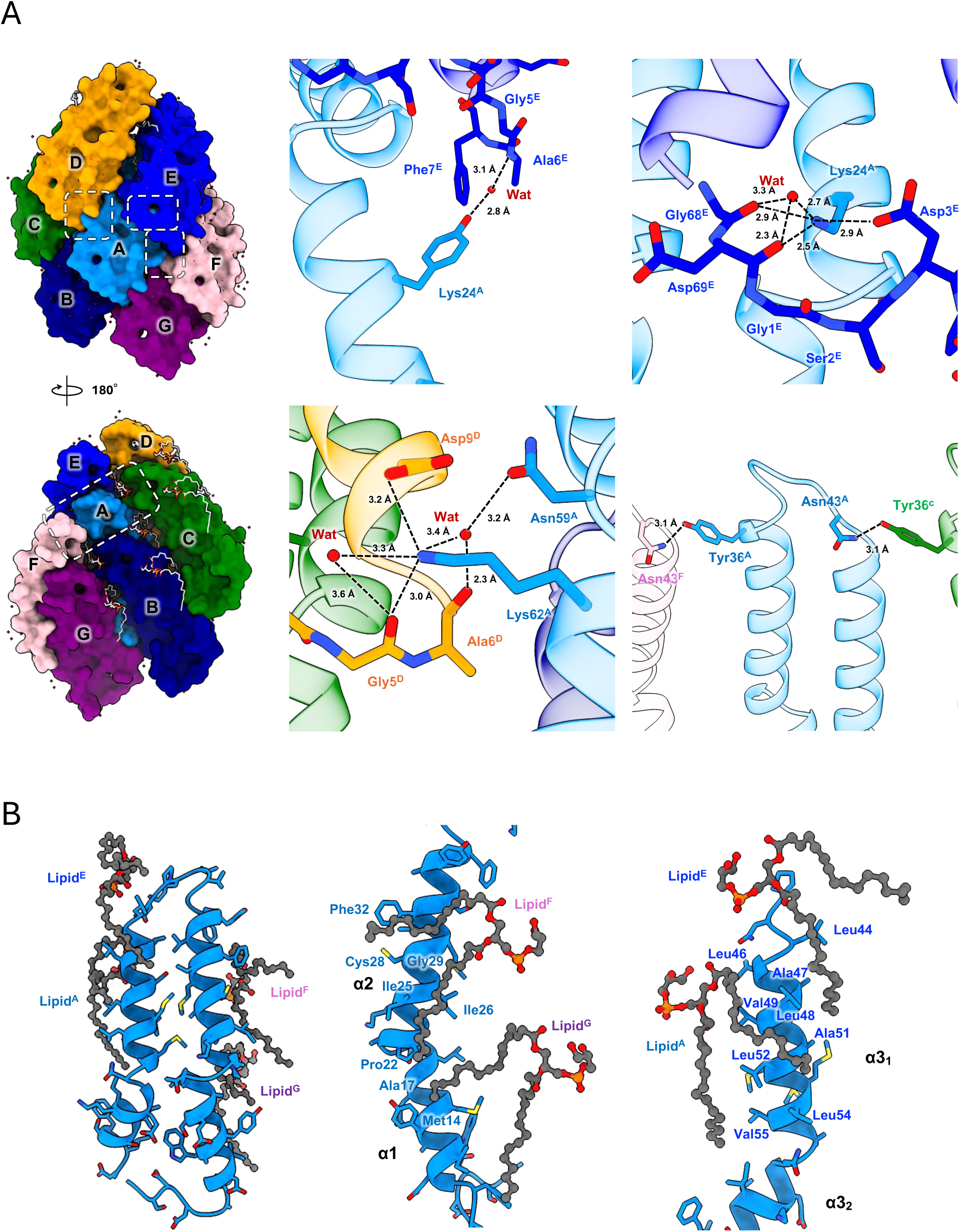
Network of interactions. (A) Extensive intermolecular and intramolecular water-mediated interactions stabilize the pilus assembly. Dashed boxes indicate key interaction interfaces. (B) The acyl chains of neighbouring PG lipids provide further stability to the H-pilus. The TrhA pilin backbone is shown as blue ribbons, and bound lipid molecules are represented as dim grey stick models. Key amino acid residues involved in lipid interactions are shown as sticks.

### Interaction between pilin subunits

The structural analysis reveals that each TrhA pilin establishes interactions with six neighboring subunits through an intricate network of both non-specific hydrophobic and specific polar contacts (Fig. 5A). The resolved water molecules appear to play an important role in mediating these subunit interactions through specific hydrogen bonding networks (Fig. 5A). A notable example is the hydrogen bond formation between Tyr13 and a water molecule, which subsequently forms hydrogen bonds with the backbone of Ala6 in the adjacent subunit (Fig. 5A). Additionally, charged residues contribute significantly to subunit stabilization, with Lys24 and Lys62 in one subunit forming salt bridges with Asp3 and Asp9, respectively, in an adjacent subunit while also making hydrogen bonds with water molecules and main chain atoms (Fig. 5A). In the cyclization region, the main chain atoms of Gly1 and Asp69 participate in a hydrogen bonding network involving Lys24 and ordered water molecules, further stabilizing their structure (Fig. 5A).

Our structural analysis revealed the presence of one tightly bound phospholipid per TrhA subunit. Each subunit interacts with four lipid molecules, primarily through interactions involving hydrophobic residues (Fig. 5B; Supp. Fig. 4). The phosphate head groups of these lipids are oriented towards the pilus lumen and positioned within hydrophilic regions (Fig. 5B; Supp. Fig. 4). This specific arrangement of lipid-protein interactions appears to contribute significantly to both the structural stability of individual subunits and the overall integrity of the assembled pilus structure. These extensive networks of protein-protein and protein-lipid interactions provide a strong molecular framework that maintains the overall H-pilus structure.

## Discussion

The R27 plasmid is a prototype conjugative plasmid, which has been extensively used to model dissemination of IncH plasmids carrying resistance genes among Gram-negative bacterial pathogen. In this study we characterized the R27-encoded H-pilus. Negative staining EM revealed that the H-pilus forms long polymerized filaments, as previously reported (25), similar to the pilus encoded by plasmids belonging to diverse incompatibility groups (e.g. IncF, IncP, IncW) (26). However, unlike previously reported conjugation pilus structures our high resolution cryo-EM structure revealed that the TrhA pilin is cyclic, with the N- and C-termini covalently bound to each other. Our mass spectrometry data confirm that cyclization is accompanied by cleavage of the last five C-terminal residues. Cyclization is driven by condensation of the Gly1 amine and Asn69 carboxylate residues. Pilin cyclization has been reported in earlier studies of the P-pilus from the IncP-1α plasmid RP4, where cyclization of the pilin TrbC proceeds via a similar condensation step as we found for the H-pilus (26); the cyclization of TrbC was proposed based on mass spectrometry and mutagenesis data. Similar to the TrhA pilin, where the last 5 terminal residues are proteolytically cleaved, 27 terminal residues of TrbC are also cleaved, although the terminal residues of the two pilins do not share any sequence similarity. It was suggested that the RP4-encoded signal peptidase homologue TraF attacks the C-terminus of TrbC via an activated serine residue. Similarly, the R27 plasmid contains TrhF, an RP4-TraF homologue (27); therefore, the mechanism of cyclisation of the H-pilus may be similar to that proposed for the P pilus.

While the IncH and IncF plasmid encode an MPFF conjugation machinery type (19), only the H-pilin is cyclic. It is widely believed that following pilus retraction both IncH and IncF plasmids use MPS, mediated by the plasmid-encoded TraN. We have previously shown the presence of at least four distinct TraN isotypes amongst IncF plasmids, each of which binds a distinct OMP (5), (6). Interestingly, TraN encoded by IncH plasmids is about 50% longer than its IncF homologues. At present, it is not known if there is a functional relationship between expression of a long TraN and the cyclic nature of the H-pilus pilin.

Cyclization has also been reported for the *Agrobacterium tumefaciens* T pilus pilin VirB2 based largely upon mass spectrometry (26, 28, 29). However, three recent cryo-EM structures of the T-pilus disagree with the MS, showing that VirB2 pilin is acyclic (24),(17),(30). It should be considered that the different conclusions might be due to different growth conditions and sample preparation used in these studies. N-terminal to C-terminal cyclization is common in peptides but rare in proteins (31, 32) and is usually associated with extreme stability under harsh conditions. Most notable cyclized structures are those of antibiotics (eg. bacitracin)(33), bacteriocins (eg. enterocin AS-48)(34), and cyclotides (e.g., Kalata B1)(35). Kalata B1 is a cyclic peptide from the cyclotide family, from the plant *Oldenlandia affinis*, known for its exceptional stability due to its cyclic cystine-knot structure (36). The cyclization of Kalata B1 occurs between Gly1 and Asn29 (35), similar to the TrhA cyclization between Gly1 and Asn69. Although cyclization of the TrhA pilin does not affect the overall H-pilus filament structure, it is very likely to provide it with additional stability under harsh conjugation conditions. This assumption is further supported by the orientation of the side chain of Lys24 that points towards the ring-like structure and provides additional hydrogen-bonds.

In summary, we have shown both structurally and biochemically that TrhA is a cyclic pilin. While mass spectrometry suggested that the P-pilus is also composed of a cyclic pilin, this has not yet been shown structurally. Mass spectrometry did suggest that the T-pilus is cyclic, but structural data did not support the cyclic nature of the T-pilin. In contrast, we show the structure of a conjugation pilus composed of a cyclic pilin. These data raise intriguing questions about the biological consequences of a cyclic pilin and the cyclization mechanism.

## Materials and Methods

### Generating MG1655 Δ*trp* and Δ*fimA* mutants

All genome editing was conducted by two-step seamless markerless homologous recombination as previous described (37), (38) (Supplementary Table 1). Briefly, mutagenesis constructs were generated by polymerase chain reaction (Q5 polymerase, New England Biolabs, UK) and Gibson Assembly (New England Biolabs, UK) on a pSEVA612S background maintained in from *E. coli* CC118 λ pir cells. Triparental mating (*E. coli* pRK2013 helper) was used to introduce mutagenesis vector into target bacterial cells, transformed with pACBSR Sm. Genomic mutations were checked by PCR and confirmed by sequencing.

### Bacterial conjugation

*E. coli* MG1655 Δ*trp* constraining the derepressed *htdA* R27 mutant (drR27) (21) was used as a donor; WT MG1655, *C. rodentium* (strain ICC169)(39) and *K. pneumoniae* (strain ICC8001)(5) were used as recipients. The donor and recipients were ground overnight in LB at 37°C, mixed in a ratio of 10:1, diluted 1 in 25 in PBS and 40 μl spotted onto LB agar plates (no selection). Following incubation at 37°C for 3 h, conjugation was allowed to proceed overnight at 25°C. The conjugation mixture was collected and resuspended in 1 mL of PBS following serial dilution from 10^-1^ to 10^-8^. The serial dilutions were plated on a M9 salts minimal media plate for selection of recipients and a chloramphenicol selection plate for selection of transconjugant cells before incubating at 37 °C for 24 h.

### Expression and purification of H-pilus

The H-pilus was purified following the protocol used to prepare the pQpKIL pilus (23). In brief, drR27 was introduced into *E. coli* MG1655 Δ*fimA*. Bacterial cultures were grown overnight in 10-20 mL LB media with Chloramphenicol (final concentration of 10 μg/mL) at 37°C. The overnight culture was placed on ice before inoculating fresh LB media. 10 mL of preculture media was used to inoculate to 1 L LB media with shaking overnight (∼14 h) at 180 rpm at 25°C. The following steps were performed at 4°C. The bacterial cells were harvested by centrifugation at 7,000 xg for 20 min and the pellet was resuspended in 20 mL PBS per 1 L of culture. The resuspended bacterial cells were shaved through 25 G needles 25∼30 times and were centrifuged at 50,000 xg for 1 h. The supernatant was collected, and PEG 6,000 was added at a final concentration of 5% with stirring for 1 h, which was followed by centrifugation at 50,000 xg for 30 min. The supernatant was decanted, and the pellet was resuspended in 200 μL imaging buffer (50 mM Tris-HCL pH 8, 200 mM NaCl). The solution containing the filaments was dialyzed overnight in imaging buffer. Sample purity was judged by SDS-PAGE.

### Negative stain EM

Purified pilus at 0.01mg/mL was applied to a glow-discharged carbon grid (300 Mesh Grids Copper). Any excess liquid was blotted off using a clean filter paper. This step was repeated by adding 3 µL of distilled water to the grid, followed by blotting. Afterward, 3 µL of 2% uranyl acetate stain was applied to the grid for negative staining, and the grid was blotted again to remove any excess stain. Finally, 3 µL of distilled water was added to rinse the grid, and the final blotting step was performed. A similar protocol was used for the preparation of cells expressing the H-pilus. The grids were imaged on a Transmission Electron Microscope (JEM-2100).

### Grid preparation and cryo-EM data collection

For grid preparation, 3 µL of the purified sample at ∼4 mg/mL was applied to a glow-discharged carbon grid (Quantifoil R1.2/1.3, Cu, 300 mesh). The grid was blotted with a blotting force of 0 for 3 sec at 4°C and 100% humidity, and flash-frozen in liquid ethane using a Vitrobot Mark IV (Thermo Fisher Scientific). The frozen grid was stored in liquid nitrogen until data collection. Cryo-EM data were collected on 300 kV Titan Krios G2 (Thermo Fisher Scientific) with a Falcon 4i detector and Selectris X energy filter at the Electron Bioimaging Center (eBIC), UK. A total of 869 movies from two grids were collected at a magnification of ×130,000 (pixel size of 0.921 Å/pixel) with a total electron dose of 50 e/Å2. The defocus range was set between −0.8 and −1.8 µm. All data were automatically acquired using EPU software. The data collection parameters are summarized in Supplementary Table 2.

### Cryo-EM Data Processing and Model Building

The collected data were processed using cryoSPARC (v.4.6.1) (22). Data were collected from two grids, analyzed separately, and the particle sets were merged for reconstruction. Both datasets were processed following similar steps. Motion correction was performed using Patch Motion Correction, and CTF estimation for micrographs was done by Patch CTF estimation (22). Particles were automatically picked using a Filament tracer without templates and extracted in a binned state at 2.76 Å/pix. The extracted particles were classified in 2D, and selected high-quality particles were used as references for the Filament tracer. 2D classification of particles from the whole dataset was used to select particles for helical reconstruction. We generated the average power spectrum using particles from the best 2D class average for the indexing of helical symmetry parameters. Helical indexing from the power spectrum suggested an axial rise of ∼12.1 Å and a twist of ∼28.9⁰, along with 5-fold point group symmetry. These symmetry parameters were used as starting parameters for helical reconstruction using Helix refine. The final H-pilus helical reconstruction showed 2.2 Å resolution using 123,942 particles, with a rise of 12.14 Å and a twist of 28.93°.

*Ab initio* model building and real space refinement were performed in PHENIX (40), (41). The model was subjected to iterative cycles of refinement and manual rebuilding in COOT (42). The structural model was validated using MolProbity (43). The processing parameters and refinement statistics are summarized in Supplementary Table 2.

### Mass spectrometry analysis

The intact mass for the protein samples were recorded on a LC-MS Waters Xevo G2 ToF MS coupled to a ACQUITY UPLC I-Class LC system (Waters Corporation). The spectrometer was auto-calibrated throughout the experiment via an internal lock-spray mass calibrant (at 556.2771 m/z of leucine enkephalin, 1 s scan every 30 s interval). The data was acquired in Continuum format, in a 500 – 2500 m/z spectral window with a scan time of 1 s and an interscan time of 0.014 s. The mass spectrometer and ESI ionization source operated under following parameters: capillary voltage 3 kV, cone voltage 40 V, source offset voltage 40 V, source temperature 100 °C, desolvation temperature 250 °C and desolvation gas (nitrogen) flow 600 L/h, cone gas flow 50 L/h. In a typical analysis, 1 µL (30 ng) of the protein sample was injected and desalted inline on a MassPrep micro desalting cartridge column (5mm x 2.1 mm 20 μm particle size, Waters™) column maintained at 60 oC. The elution was performed using solvent A (95% water, 5% acetonitrile with 1% formic acid) and solvent B (95% acetonitrile 5% water with 1% formic acid) with a gradient elution using isocratic 98% solvent A (0 – 0.5 min), linear gradient to 98% solvent B (0.5 – 3.8 min), and isocratic 98% solvent B (3.8-4.5 min) before return to 98%A (4.5-4.6 min) and isocratic at 98%A (4.6-5min) at a total flow of 0.2 ml/min. Combined mass spectra from the total chromatographic protein peak were used for intact mass reconstruction using MassLynx (Waters) Max(imum) Ent(ropy) 1 deconvolution algorithm (Resolution: 1.00 Da/channel, Peak Width at half height of 0.1, Minimum intensity ratios: 33% Left and Right). Spectra were deconvoluted between 6500 – 8000 Da.

## Data availability

The cryo-EM maps have been deposited in the Electron Microscopy Data Bank (EMDB) under accession code EMD-52431. The structural coordinates have been deposited to the RCSB Protein Data Bank (PDB) under the accession code 9HVC. Raw images were submitted to the Electron Microscopy Public Image Archive (EMPIAR) with ID xxxx.

## Acknowledgements

We would like to acknowledge Diamond for access and support of the cryo-EM facilities at the UK national electron Bio-Imaging Centre (eBIC). NI is funded by a Fellowship from The Naito Foundation. CS was funded by a BBSRC DTP Studentship grant (BB/M011178/1). EHE was funded by NIH GM122510.

## Author contributions

KB and GF conceived and supervised the study. NI and CS purified the H-pilus. NI determined the cryo-EM structure. NI and RS performed model building and refinement. NI and NS imaged the negative stained grids. JLCW, SH performed conjugation experiments. CB isolated the R27 plasmid. OWP and AC generated the strain to purify the H-pilus. KB, GF, EE, NI and RS analyzed the structures. SS performed mass spectrometry analysis. NI, EE, GF and KB wrote the manuscript with the help of all authors.

---

**Supplementary Table 1.**
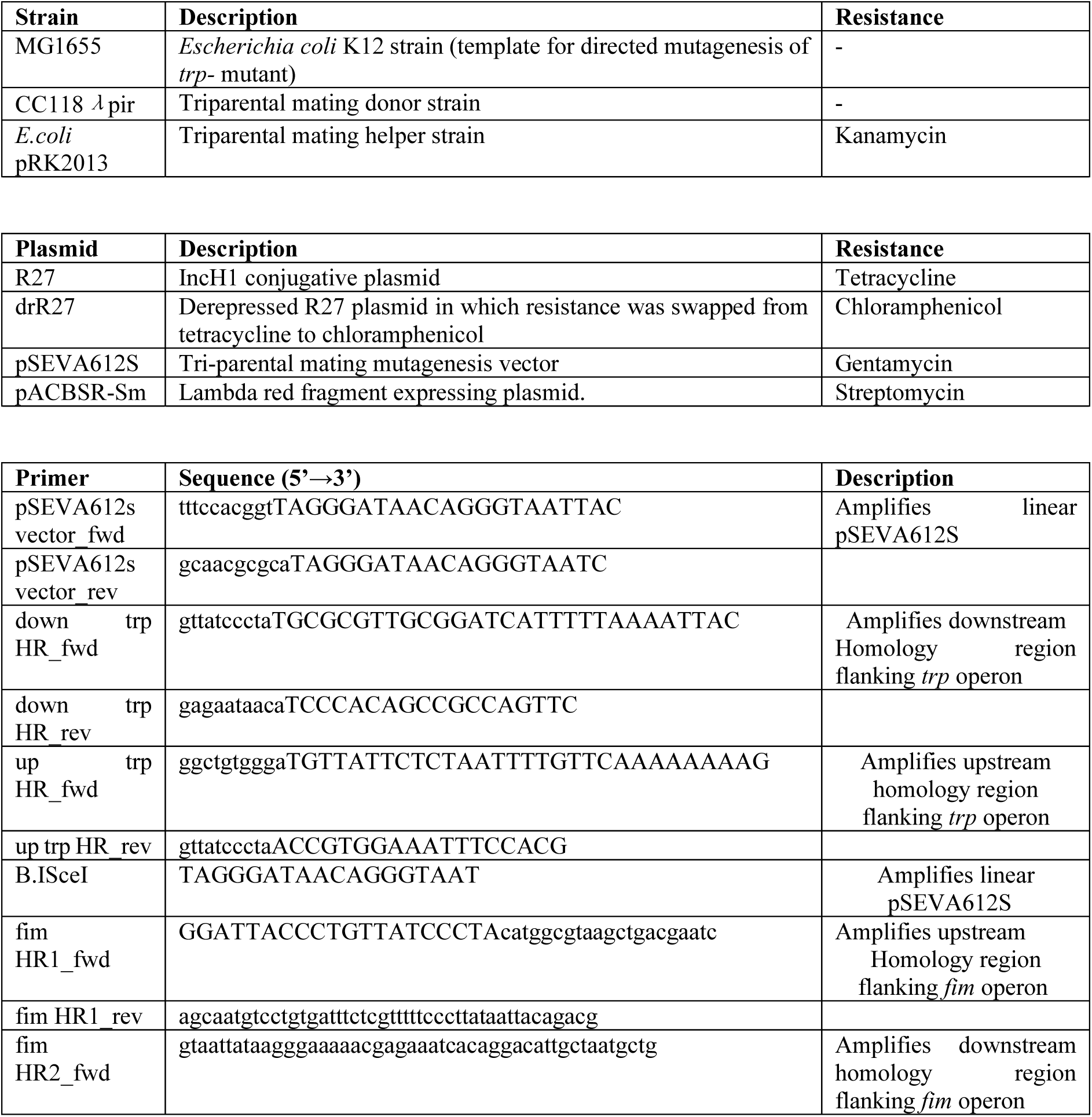
Strains, plasmids and primers used in the generation of the MG1655 Δ*trp* and Δ*fimA* mutants.

**Supplementary Table 2.** Data collection and processing parameters, and refinement statistics.

|  |  |
| --- | --- |
|  | H-pilus |
| **PDB entry** | 9HVC |
| **EMDB entry** | EMD-52431 |
| **Data collection and processing** |  |
| Magnification | 130,000 |
| Microscope | Titan Krios G2 |
| Voltage (kV) | 300 |
| Detector | Falcon 4i |
| Electric exposure (e<sup>-</sup>/Å) | 50 |
| Defocus range (μm) | -0.8 to -1.8 |
| Pixel size (Å) | 0.921 |
| Data Processing Program | cryoSPARC (v.4.6.1) |
| Movies | 869 |
| Initial / Final particle images (no.) | 492,091 / 123,942 |
| Symmetry imposed | C5 |
| Helical rise (Å) | 12.14 |
| Helical twist (°) | 28.93 |
| Map resolution (Å) | 2.2 |
| FSC threshold | 0.143 |
| **Refinement** |  |
| Refinement Program | PHENIX (v.1.20.1) |
| Map:model resolution (Å) | 2.2 |
| FSC threshold | 0.5 |
| Model composition |  |
| Non-hydrogen atoms | 578 |
| Protein residues | 69 |
| Water | 31 |
| R.m.s. deviations |  |
| Bond length (Å) | 0.003 |
| Bond angles (°) | 0.388 |
| Validation |  |
| MolProbity score | 1.22 |
| Clashscore | 4.45 |
| Ramachandran plot |  |
| Favored / Allowed (%) | 100 / 0 |
| Disallowed (%) | 0.00 |
| Mask CC | 0.88 |

**Supplementary Figure 1.**
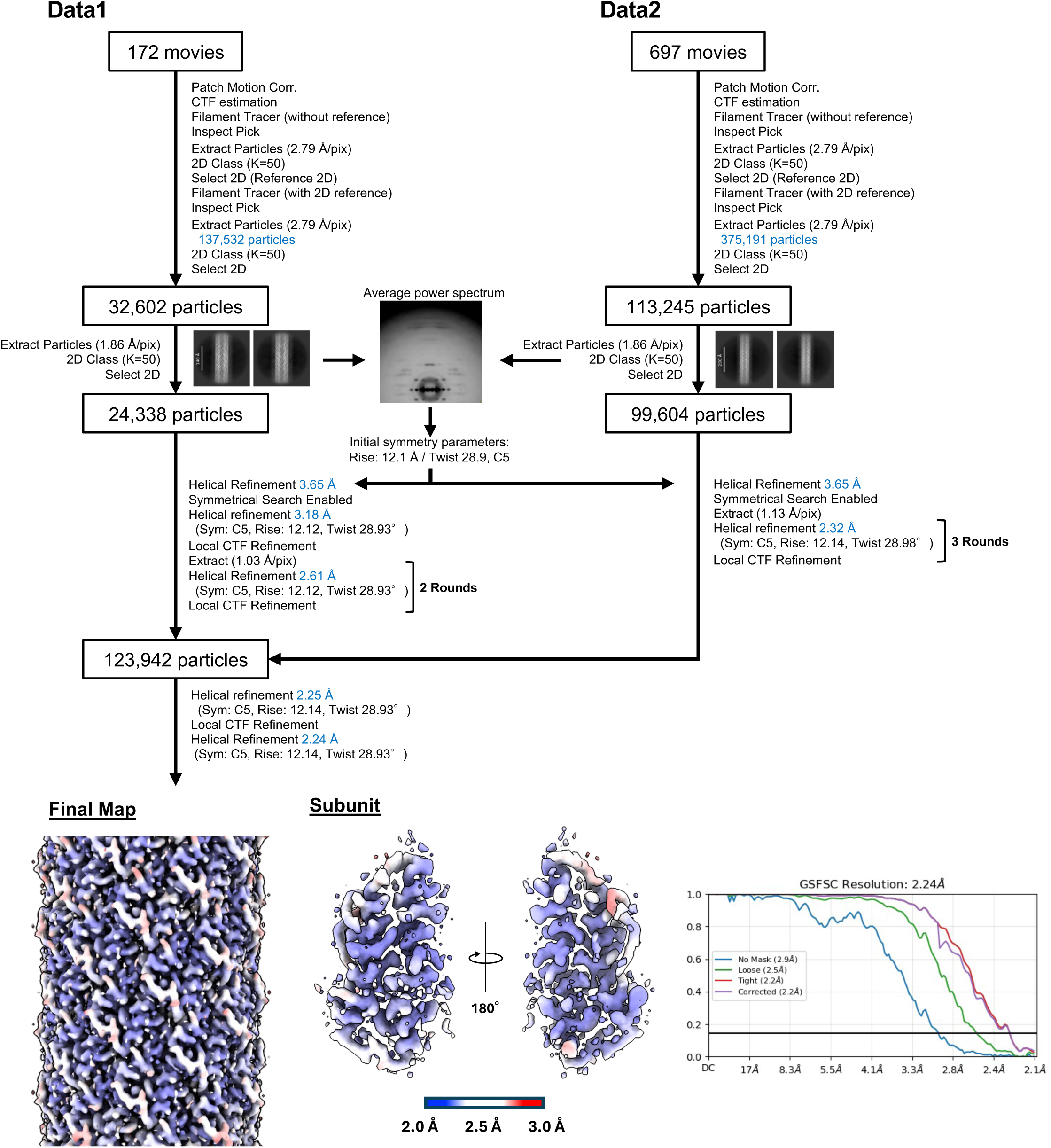
Cryo-EM data processing workflow. Data processing workflow for H-pilus. All the processing was performed in cryoSPARC (v.4.6). The Final map was colored according to the local resolution. Gold-standard FSC curve of the final map is shown. The resolution was determined at FSC=0.143.

**Supplementary Figure 2.**
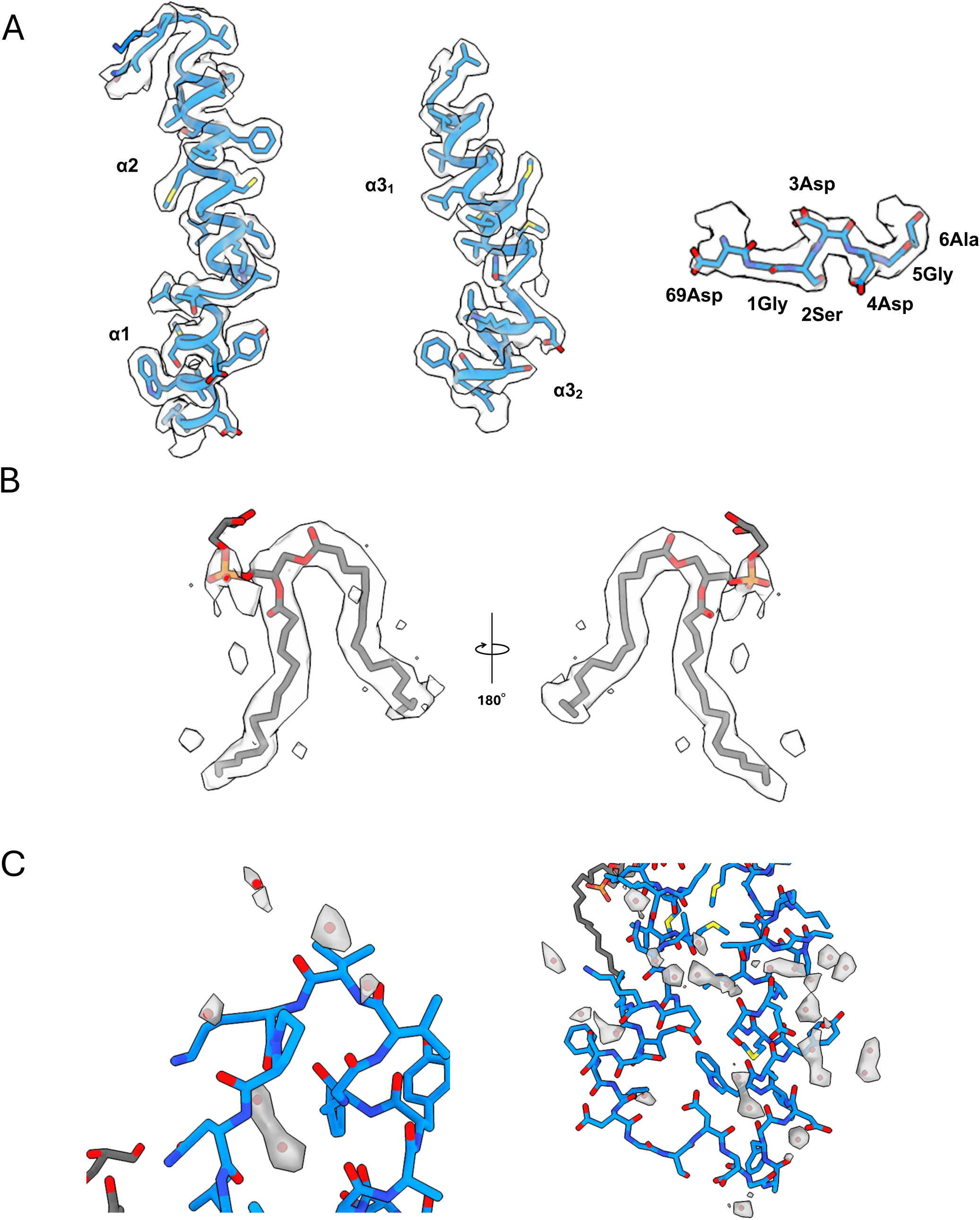
Cryo-EM density map and models. The map and models showing: (A) helices α1,2,3 and cyclization region; (B) lipid; and (C) water molecules.

**Supplementary Figure 3.**
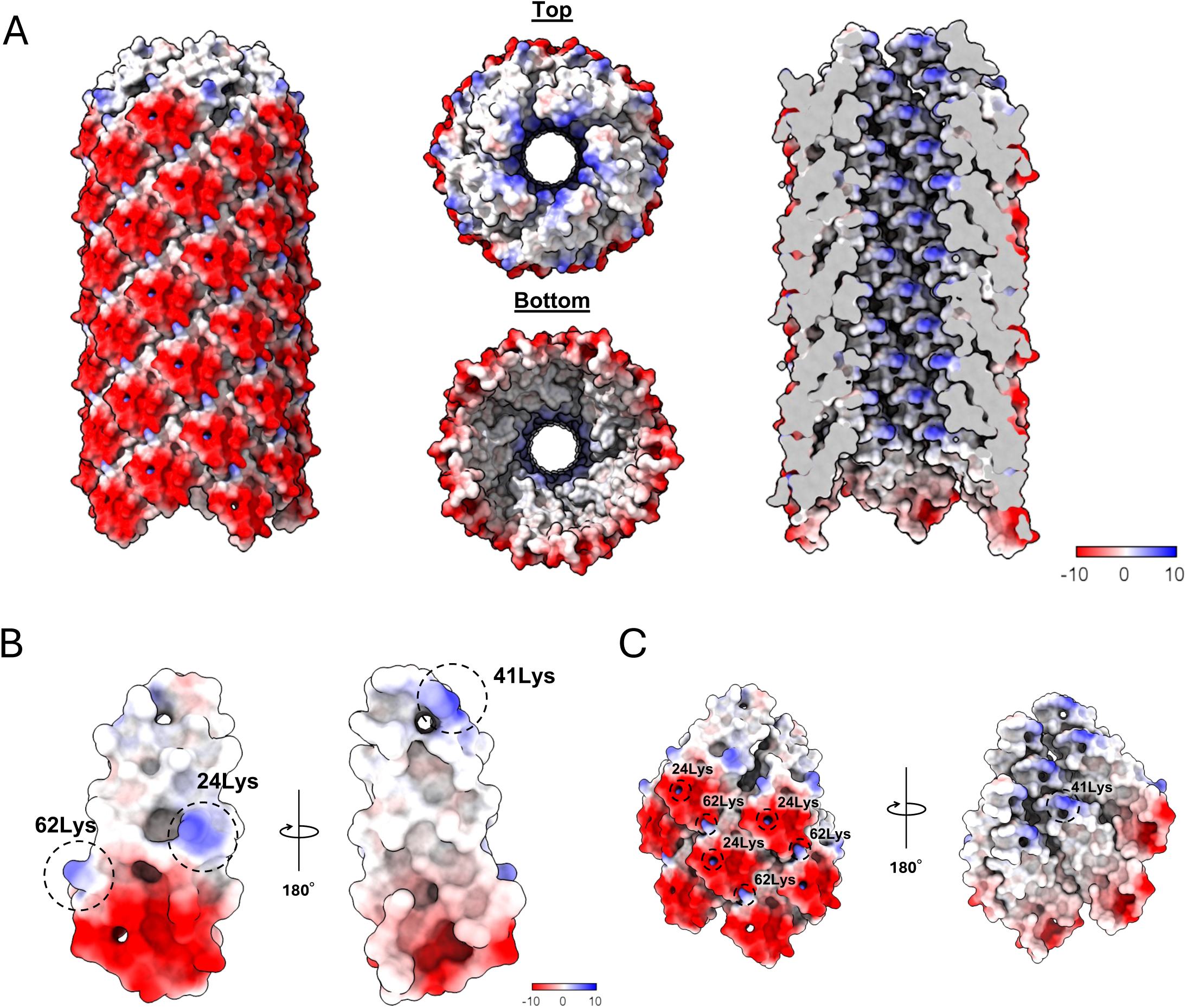
The surface electrostatic potential of H-pilus. All potentials have been calculated without the bound lipid. (A) The potential is shown in the side view (left), top and bottom views (middle), and cutaway view (right). The electrostatic potential is colored from red (negative) to white (neutral) to blue (positive). (B) A detailed surface electrostatic surface of the TrhA pilin is shown in two orientations. Key lysine residues (Lys24, Lys41, and Lys62) are highlighted with dashed circles. (C) Residues in neighboring subunits are labelled, and analysis of electrostatic interactions between subunit and subunit. The view is shown in two orientations, highlighting key lysine residues.

**Supplementary Figure 4.**
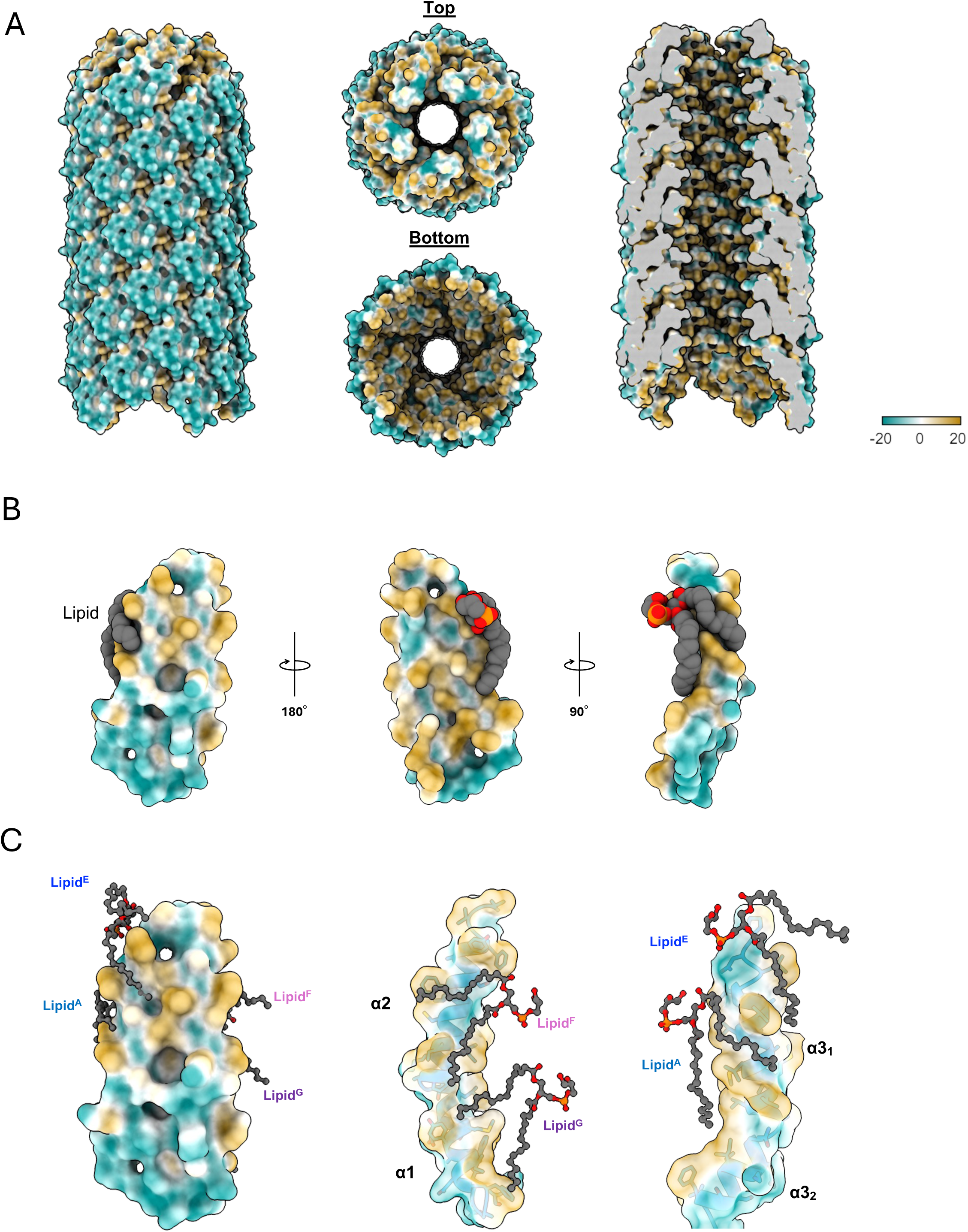
The surface hydrophobicity analysis of H-pilus. (A) The surface hydrophobicity is shown in side view (left), top and bottom views (middle), and cutaway view (right). The hydrophobic potential is colored from gold (hydrophobic) to white (neutral) to cyan (hydrophilic). (B) Surface hydrophobicity of the TrhA pilin and its association with the PG lipid is shown in three orientations. (C) Surface hydrophobicity of lipid-binding pockets in α1, α2, and α3.

